# Characterization of Viroplasm-Like Structures by Co-Expression of NSP5 and NSP2 Across Rotavirus Species A to J

**DOI:** 10.1101/2024.06.04.597348

**Authors:** Melissa Lee, Ariana Cosic, Kurt Tobler, Claudio Aguilar, Cornel Fraefel, Catherine Eichwald

**Author notes:** Address correspondence to Catherine Eichwald.

## Abstract

Rotaviruses (RV) are classified into nine species, A-C and D-J, with species A being the most studied. In rotavirus of species A (RVA), replication occurs in viroplasms, which are cytosolic globular inclusions primarily composed of the proteins NSP5, NSP2, and VP2. The co-expression of NSP5 with either NSP2 or VP2 leads to the formation of viroplasm-like structures (VLS). Although morphologically identical to viroplasms, VLSs cannot replicate, but they serve as excellent simplified tools for studying complex viroplasms.

There is a knowledge gap regarding viroplasms of non-RVA species due to a lack of research tools, such as specific antibodies and tissue culture systems. In this study, we explored the ability of NSP5 and NSP2 from non-RVA species to form VLSs. The co-expression of these two proteins led to globular VLSs in RV species A, B, D, F, G, and I, while RVC formed filamentous VLSs. The co-expression of NSP5 and NSP2 of RV species H and J did not result in VLS formation.

Interestingly, NSP5 of all RV species self-oligomerizes, with the ordered C-terminal region, termed the tail, being necessary for self-oligomerization of RV species A-C and G-J. Except for NSP5 from species J, all NSP5 bound with their respective NSP2. We also found that interspecies VLS are formed between closely related RV species B with G and D with F. Additionally, VLS from RVH and RVJ formed when the tail of NSP5 RVH and RVJ was replaced by the tail of NSP5 from RVA and co-expressed with their respective NSP2.

**Importance:** Rotaviruses (RV) are classified into nine species, A-D and F-J, infecting mammals and birds. Due to the lack of research tools, all cumulative knowledge on RV replication is based on RV species A (RVA). The RV replication compartments are globular cytosolic structures named viroplasms, which have only been identified in RV species A. In this study, we examined the formation of viroplasm-like structures (VLS) by the expression of NSP5 with NSP2 across RV species A to J. Globular VLSs formed for RV species A, B, D, F, G, and I, while RV species C formed filamentous structures. The RV species H and J did not form VLS with NSP5 and NSP2. Similar to RVA, NSP5 self-oligomerizes in all RV species, which is a requirement for VLS formation. This study provides basic knowledge of the non-RVA replication mechanisms, which could help develop strategies to halt virus infection across RV species.

## Introduction

Rotavirus (RV) is an etiological agent responsible for severe gastroenteritis in infants and young children killing approximately 128,000 children per year, mainly in low- and middle-income countries (1). Additionally, RV outbreaks are the leading cause of diarrhea among the adult population in several countries (2–6). In the US, RV infection ranks as the second most common cause of diarrhea in adults after norovirus (7–9). From a veterinary perspective, RV infections significantly impact livestock worldwide. RV accounts for 80% of diarrhea cases in piglets in the USA, Canada, and Mexico, with potential zoonotic implications in humans (10). In the poultry industry, RV infections impact the feed conversion ratio, resulting in substantial economic losses (11).

RV of species A (RVA) is a segmented dsRNA virus belonging to the *Reoviridae* family. The RV virion is organized in three concentric layers surrounding the viral genome. The spike protein VP4 and the glycoprotein VP7 compose the outermost layer, while VP6 forms the intermediate layer. The innermost layer, the core shell, comprises 120 copies of VP2, organized in twelve asymmetric decamers (12). Each core-shell encapsidates the eleven single copies of the genome segment and the replication complexes composed of the RNA-dependent RNA polymerase (RdRp) VP1 and the guanylylmethyltransferase VP3, which localize beneath each of the five-fold axes of the VP2 decamer. The external layer is lost during virus entry, and a transcriptionally active double-layered particle (DLP) is released into the cytosol (13). The newly released transcripts give rise to the viral proteins necessary for viral replication. Among those proteins, the NTPase/RTPase NSP2 and the phosphoprotein NSP5, together with the structural proteins VP1, VP2, VP3, and VP6, make part of the RV replication compartments termed viroplasms (14). The viroplasms correspond to electron-dense membrane-less globular cytosolic inclusions where viral genome transcription, replication, and the packaging of the newly synthesized pre-genomic RNA segments into the viral cores occur. The viroplasms are highly dynamic, being able to coalesce between them and move to the juxtanuclear region of the cell at increasing times post-infection (15–17). Despite not yet being well-defined, several host factors have been identified as necessary for viroplasm formation and maintenance (18–21). The initiation process for viroplasm formation requires a scaffold of lipid droplets by incorporating perilipin-1 (22, 23). Furthermore, the host cytoskeleton, actin filaments, and microtubules (MT) play a role in the formation, maintenance, and dynamics of the viroplasms (16, 24, 25). In this context, NSP2 octamers are directly associated with MTs to promote viroplasm coalescence (16, 26–29). Moreover, VP2 plays a role in viroplasm dynamics by allowing their perinuclear motion (16). Finally, consistent with these features, the viroplasms have been attributed to liquid-liquid phase-separated structures (30). Interestingly, co-expression of the main viroplasm protein NSP5 with either NSP2 or VP2 leads to the formation of cytosolic inclusions named viroplasm-like structures (VLS), which are morphologically similar to viroplasms but unable to yield viral progeny (15, 16, 31–34).

NSP5 is required for viroplasm formation and virus replication (35–37), having a multifunctional role in the RV life cycle, interacting with NSP6 (34), NSP2 (15), VP1 (38), VP2 (39, 40), and unspecifically to dsRNA (41). These attributes are consistent with its predicted unfolded nature (42–44). Interestingly, the C-terminal region of NSP5 is needed for its self-oligomerization (34, 45), to associate with other RVA proteins (15, 34, 38, 40), and to form the viroplasms (37). NSP5 is sumoylated (46), presumably a pre-requirement for interacting with viral or host components. NSP5 is also phosphorylated, which is crucial for viroplasm morphology (37), a trait for liquid-liquid phase separation conditions of the viroplasms (30). NSP5 hyperphosphorylation is triggered by the association with NSP2 or VP2, primed at serine-67 by casein kinase 1 alpha (31, 45, 47, 48). Collectively, NSP5 is a crucial component in RVA replication.

NSP2 self-assembles in a 4-2-2 crystal symmetry consisting of two doughnut-shaped tetramers with a highly electropositive nature (27, 49). NSP2 is linked to several enzymatic activities such as nucleoside diphosphate kinase-like (50), RNA-helix-destabilizing (50), and nucleoside triphosphatase activities (27) consistent with molecular motor properties. Also, NSP2 phosphorylation has been implicated in viroplasm formation, as evidenced by the delayed formation of viroplasms observed in an NSP2 S313D phosphomimetic mutant. NSP2 directly influences viroplasm coalescence (15) through association with MTs (28). The flexible NSP2 C-terminus also improves viroplasm morphology (51) and its activity as an RNA chaperonin (26). Interestingly, NSP2 binds to VP1 and viral RNA (52, 53), which implicated it as a factor in replication intermediates in the viroplasms.

All the current information compiled on the RV replication life cycle is based on species A, which has a broad spectrum of strains that mainly infect young mammals like infants, piglets, and calves. According to the International Committee on Taxonomy of Viruses (ICTV), RVs are currently grouped into nine species (54): A to D and F to J. These species have been detected in diverse hosts. Rotavirus B (RVB) has been identified in human adults, rats, cattle, goats, sheep, and swine (55). RVs D (RVD), F (RVF), and G (RVG) have been only detected in avian species. Studying the replication of these RV species is challenging since most of the knowledge about them has been obtained by in-depth sequencing of nucleic acid isolated from infected samples, and therefore, no virus inoculum is available for their investigation. The few isolated viruses of non-RVA species are not well adapted to tissue culture (56), and research tools such as specific antibodies are unavailable. Therefore, all information about RV replication and life cycle corresponds to RVA species. In this context, there is no evidence in non-RVA species of viroplasm formation, which is crucial for virus replication. A comprehensive approach to the mechanism of replication of non-RVA species is essential for tackling RV infection in diverse hosts, including humans (RVB), pigs (RVA and RVC), and avians (RVD, RVF, and RVG). RV reverse genetics has been recently established only for some RVA strains like simian SA11(57), porcine OSU(58), and human KU(59). Therefore, using this technology for other RV species is still unfeasible.

In this study, we investigated whether rotavirus (RV) species B to J can form globular viroplasm-like structures (VLS) by co-expressing their corresponding NSP5 and NSP2 proteins. We demonstrate that NSP5 in all tested RV species can self-oligomerize, primarily through its predicted structurally ordered region. Additionally, we tested the ability of NSP5 to bind NSP2 in non-RVA species. Finally, we examined RV interspecies formation of VLS and found that some RV species can form heterologous NSP5- and NSP2-induced VLS.

## Results

### NSP5 and NSP2 across species A to J

Viroplasms, globular cytosolic replication compartments, have been described exclusively for RVA. No evidence points to the presence of these structures in other RV species. Interestingly, the co-expression of two RVA proteins, NSP5 and NSP2, can lead to the spontaneous formation of VLS, which are morphologically identical to viroplasms but unable to yield virus progeny. VLSs are excellent tools for studying the molecular details of the formation of viroplasms. In this context and in the absence of cell culture systems that allow investigating the replication of non-species A RVs, we assessed if NSP5 and NSP2 of RV species B to J can form globular VLSs comparable to the ones observed in RVA. For this purpose, we identified in the NCBI data bank couples of NSP5 and NSP2 of RV species B to J (**Table 1**). As a reference for RVA, we used the ORFs of NSP5 and NSP2 of simian strain SA11. The sequence alignment (**Fig S1**) for NSP5 and NSP2 found poor similarity across the RV species, particularly for NSP5. Even though these proteins have an equivalent number of residues, ranging from 157 (RVI) to 218 (RVF) for NSP5 and from 296 (RVG) to 318(RVF) for NSP2, they are quite diverse. Similar to NSP5 RVA (60) (**Table 1**), NSP5 of the other RV species has a high content of serines and threonines, ranging from 14.37% for RVG to 23.74 % for RVA. NSP5 RVA has been demonstrated to be an intrinsically disordered protein (IDR) with an ordered C-terminal region of approximately 18 amino acids (30). In fact (**Fig 1a**), this can be visualized using a PONDR score (http://www.pondr.com) that compares the primary amino acid sequences of NSP5 RVA from simian strain SA11 and porcine strain OSU. The PONDR score analysis showed the same disordered arrangement for most NSP5 RV species (**Fig 1b**), except for NSP5 RVC, RVD, and RVF (**Figs 1c-e**). Thus, NSP5 of RVC has central and C-terminal regions that are predicted to be ordered. Meanwhile, NSP5 of RVD and RVF have a predicted ordered N-terminal region. Consistent with the PONDR score, AlphaFold3 predicted an alpha helix at the C-terminal region of the monomeric NSP5 of RVA, RVB, RVG, RVH, RVI, and RVJ (**Fig S2**). Accordingly, an alpha helix is also predicted at the C-terminal region of NSP5 from RVC. However, the prediction of the presence of an alpha helix at either N- or C-terminal regions is of reduced confidence for RVD and RVF. Moreover, for RVD an alpha helix is predicted in the NSP5 central region, between glutamic acid 73 and serine 111.

**Figure 1.**
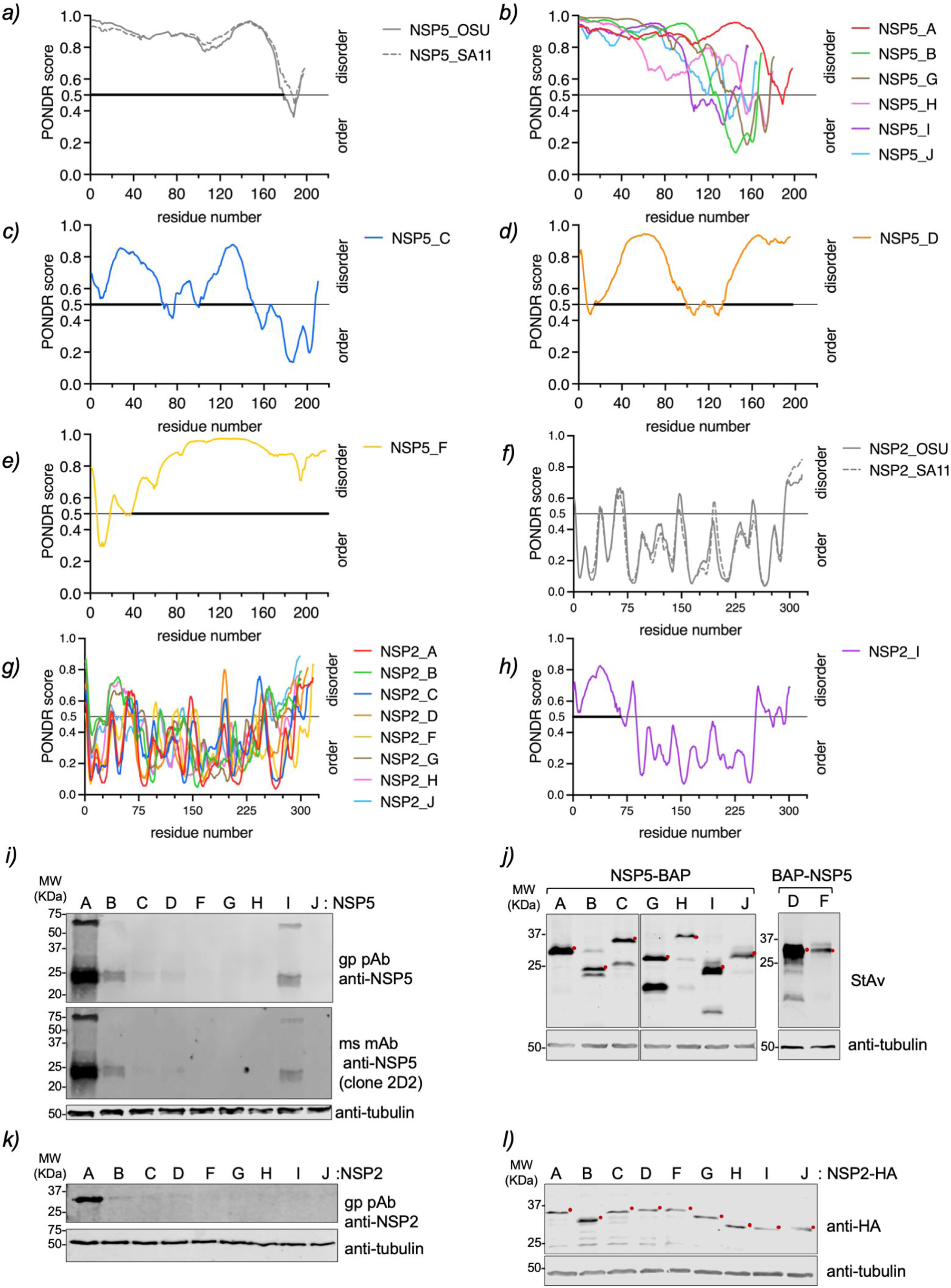
IDR and expression of NSP5 and NSP2 of species A to J. Plots comparing IDR prediction of NSP5 of RVA strains OSU and SA11 **(a)**, RVA, RVB, RVG, RVH, RVI, RVJ **(b)**, RVC **(c)**, RVD **(d)** and RVF **(e)**. Plots comparing IDR prediction of NSP2 of RVA strains OSU and SA11 **(f)**, RVA-RVH, RVG **(g),** and RVI **(h)**. The bold grey line highlights the disordered region in the protein. **i)** Immunoblotting of MA104 cells expressing NSP5 of RVA to RVJ. The membrane was incubated with guinea pig polyclonal anti-NSP5 (top panel) and mouse monoclonal antibody (mAb) anti-NSP5 clone 2D2 (middle panel). **j)** Immunoblotting of MA/cytBirA cells expressing NSP5-BAP of RVA-RVC, RVG-RVJ, and BAP-NSP5 of RVD and RVF. The membrane was incubated with streptavidin(StAv)-IRDye800. **k)** Immunoblotting of MA104 cells expressing NSP2 of RVA-RVJ. The membrane was incubated with guinea pig polyclonal anti-NSP2. **l)** Immunoblotting of MA104 cells expressing NSP2-HA of RVA to RVJ. The membrane was incubated with mouse mAb anti-HA. Anti-tubulin was used as a loading control. The red dot indicates the predicted molecular weight of the recombinant proteins.

**Table 1.**
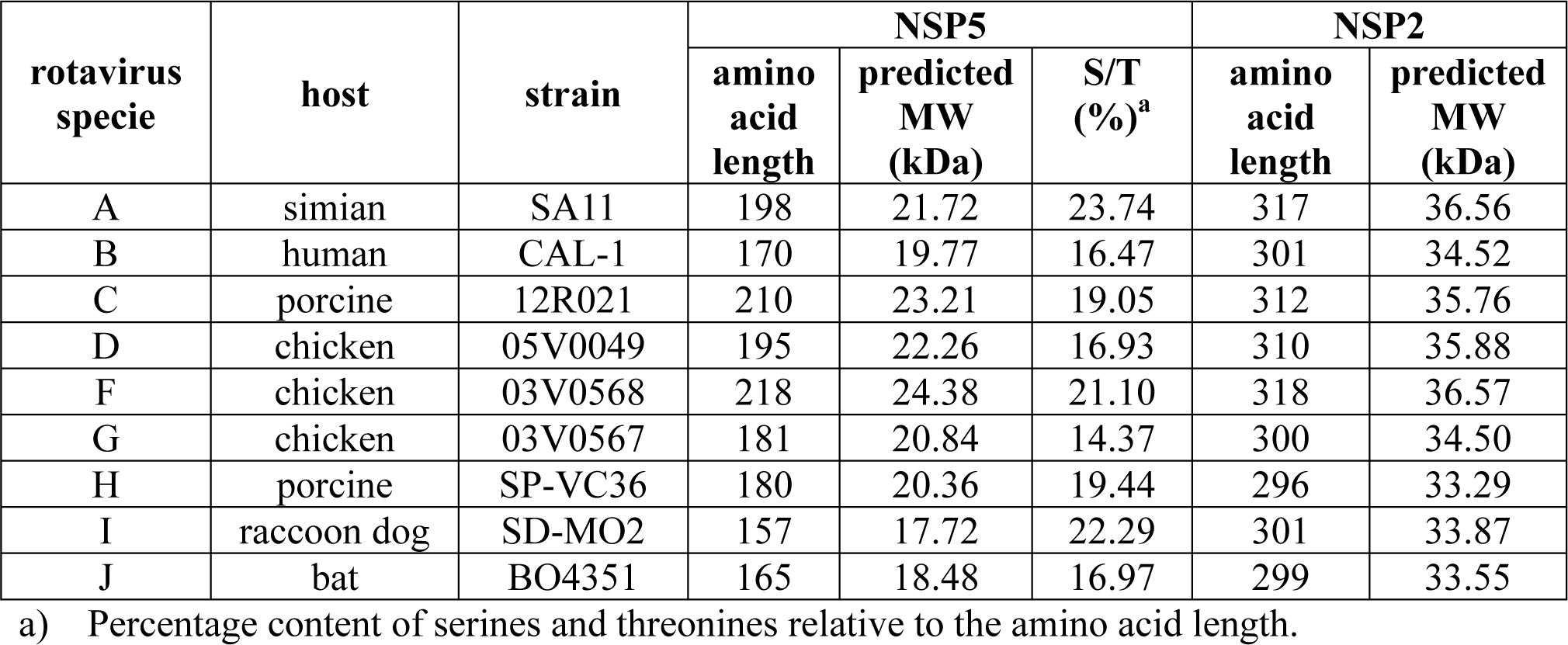
NSP5 and NSP2 protein features of the RV species analyzed in this study.

In contrast, NSP2 of RV species A, B, C, D, F, G, H, and J have similar PONDR score patterns consistent with an ordered protein (**Fig 1f and g**). The N-terminal region of NSP2 RVI is predicted to be disordered compared to NSP2 of the other RV species (**Fig 1h**). NSP2 of RVA, RVB, and RVC have already been described as forming octamers with a 4-2-2 conformation (49, 61, 62). Interestingly, AlphaFold3 (63) predicts that NSP2 forms octamers with a 4-2-2 conformation for all RVA species, A to J (**Fig S3**).

Each open reading frame of NSP5 and NSP2 of RV species B to J was chemically synthesized, cloned in expression plasmids, and tested for expression in MA104 cells by immunoblotting. As expected (**Fig 1i**), staining with guinea pig anti-NSP5 (32, 33) and mouse monoclonal anti-NSP5 (clone 2D2) antibodies (64) resulted in strong NSP5/A but only weak NSP5/B and NSP5/I signals. Similarly (**Fig 1k**), only NSP2/A was detected in immunoblotting with guinea pig anti-NSP2 antibody (48). These outcomes are consistent with the high antigenic diversity between NSP5 and NSP2 across RV species A to J. Therefore, NSP5 and NSP2 were fused to a biotin acceptor peptide (BAP) and HA tags, respectively, to enable their detection through diverse biochemical assays. Specifically, NSP5/A, B, C, G, H, I, and J were fused at the C-terminus with a BAP tag (NSP5-BAP), and NSP5/D and F were fused at the N-terminus with a BAP tag (BAP-NSP5) (**Fig 1j**). All recombinant NSP5 proteins were shown to be correctly biotinylated when expressed in MA/cytBirA cells, a cell line expressing a cytosolic biotin ligase BirA (65), since they migrated at the predicted molecular weight (**Table 2**). The NSP2-HA proteins of the diverse RV species also proved to migrate at the expected molecular weights (**Fig 1l and Table 2**).

**Table 2.** Predicted molecular weight of NSP5 and NSP2 proteins from divers RV species fused to BAP or HA.

| **RV specie** | **Host** | **Strain** | **NSP5<sup>a</sup>**<br>**(kDa)** | **NSP5-BAP<sup>a</sup>**<br>**(kDa)** | **NSP2<sup>a</sup>**<br>**(kDa)** | **NSP2-HA<sup>a</sup>**<br>**(kDa)** |
| --- | --- | --- | --- | --- | --- | --- |
| **A** | simian | SA11 | 21.72 | 23.7 | 36.56 | 37.7 |
| **B** | human | CAL-1 | 19.77 | 21.7 | 34.52 | 35.6 |
| **C** | porcine | 12R021 | 23.21 | 25.2 | 35.76 | 36.9 |
| **D** | chicken | 05V0049 | 22.26 | 24.2 | 35.88 | 37.0 |
| **F** | chicken | 03V0568 | 24.38 | 26.3 | 36.57 | 37.7 |
| **G** | chicken | 03V0567 | 20.84 | 22.8 | 34.50 | 35.6 |
| **H** | porcine | SP-VC36 | 20.36 | 22.3 | 33.29 | 34.4 |
| **I** | raccoon dog | SD-MO2 | 17.72 | 19.7 | 33.87 | 35.0 |
| **J** | bat | BO4351 | 18.48 | 20.4 | 33.55 | 34.7 |

a) Predicted molecular weight (kDa).

### Characterization of VLS formation across RV species A to J

Next, we investigated if the co-expression of cognate NSP5 and NSP2 allowed the formation of VLS, which are defined by the colocalization of the signals of these two proteins in globular cytosolic inclusions. For this purpose, cognate couples of NSP5-BAP and NSP2-HA of RV species A to J were expressed in MA/cytBirA cells. As expected (**Fig 2a**), the co-expression of NSP5-BAP with NSP2-HA of RVA produced VLSs (33). When inspecting the formation of globular cytosolic VLSs across RV species B to J, we found that RV species B, D, F, G, and I also formed globular inclusions upon co-expression of cognate NSP5-BAP with NSP2-HA. Interestingly, RVC forms a mixture of small globular inclusions and filamentous structures. We confirmed this result by co-expressing NSP5-BAP and NSP2-HA of RVC at diverse ratios (**Fig S4**) and found that the overexpression of NSP5-BAP forms filamentous structures while overexpression of NSP2-HA leads to the formation of globular VLSs. The filamentous structures dissolve upon nocodazole treatment, suggesting a dependence on the MT network. Moreover (**Fig S5**), the co-expression of NSP5-BAP and NSP2-HA of RV species H and J did not lead to VLS formation at any of the tested pCI-NSP5-BAP:pCI-NSP2-HA transfection ratios (4:1, 2:1, 1:1, 1:2, and 1:4). NSP5-BAP and NSP2-HA of avian RV species D, F, and G, behaved differently from the mammalian cohorts. In this context, BAP-NSP5/D appeared homogenously distributed in the cytosol and nucleus, while BAP-NSP5/F forms globular nuclear inclusions and NSP5-BAP/G forms aggregates in the nucleus. Interestingly, when BAP-NSP5/D, BAP-NSP5/F, or NSP5-BAP/G were co-expressed with their respective NSP2-HA, they were redistributed in the cytosol-forming VLSs. The numbers of RVG-VLSs were smaller than the ones observed for RVD and RVF VLSs. Adding a BAP tag at the C-terminus of NSP5/D or NSP5/F did not allow the formation of VLSs (data not shown). Because of the peculiar distribution pattern of avian NSP5 RV species D, F, and G, we investigated if the mammalian host environment provided by the MA104 cells could affect the distribution of these proteins. For this purpose (**Fig 2b**), we expressed NSP5-V5 or NSP2-HA of RV species D, F, and G in chicken epithelial LMH cells. Thus, V5-NSP5/D and F and NSP2-HA/D and F mainly localized homogenously in the cytosol of the chicken cells. In contrast, NSP5-V5/G formed spontaneous globular cytosolic structures. The co-expression of NSP5-V5 and NSP2-HA of RV species D, F, and G in LMH cells led to the formation of globular cytosolic inclusions, particularly large for RVD and RVF. Interestingly (**Fig 2b**), VLSs built-up of NSP5-V5 and NSP2-HA were apparently less abundant than globular cytosolic structures composed by NSP5-V5/G expressed alone.

**Figure 2.**
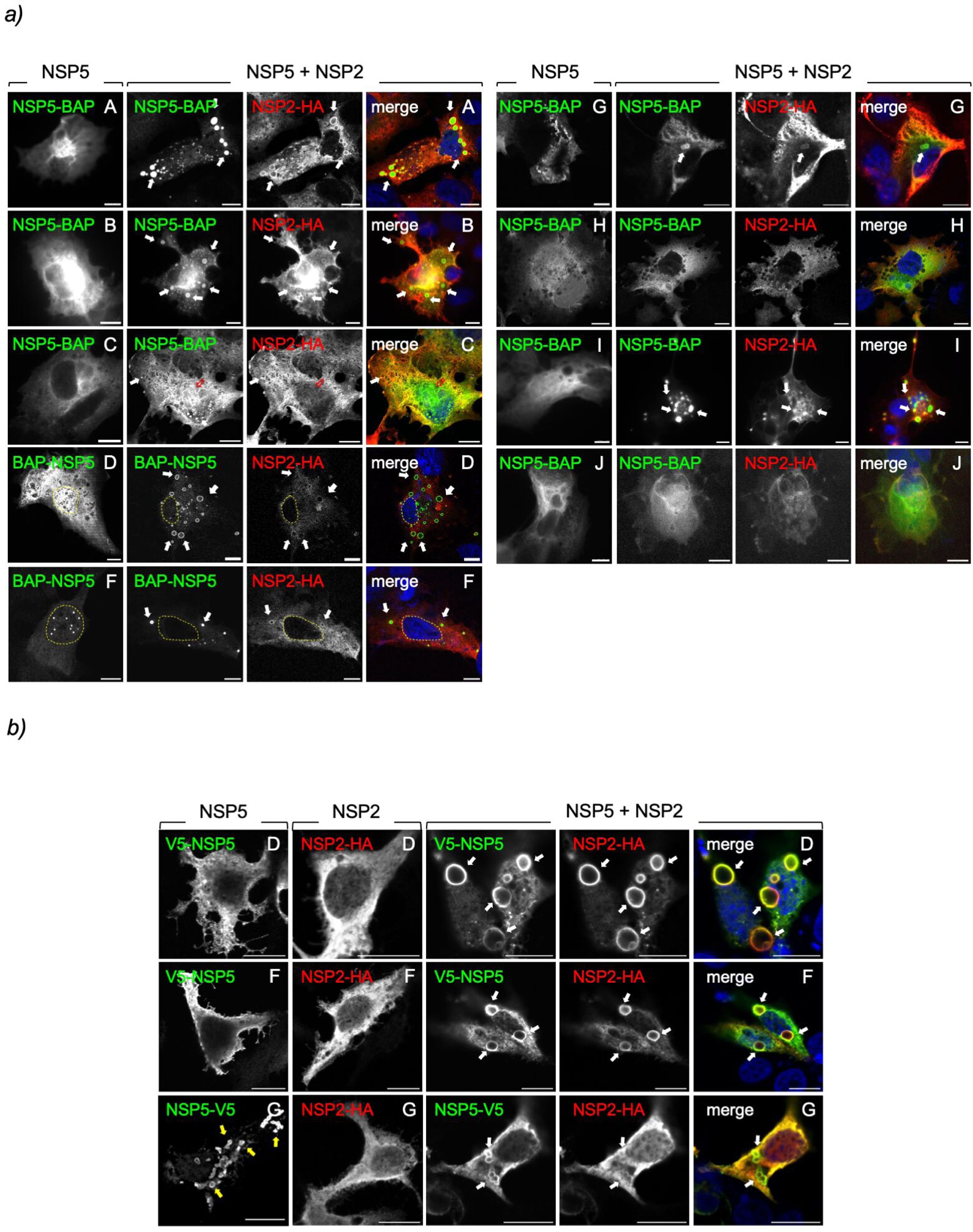
Characterization of VLS formation by co-expression of NSP5-BAP and NSP2-HA across RV species A to J. **a)** Immunofluorescence images of MA/cytBirA cells expressing NSP5-BAP (RVA-RVC and RVG-RVJ) or BAP-NSP5 (RVD and RVF) alone (left column) or in co-expression with their respective NSP2-HA. At 16 hpt, the cells were fixed and immunostained for detection of NSP5-BAP (StAv, green, first and second columns) and NSP2-HA (anti-HA, red, third column). A merged image is shown in the right column of each panel. Nuclei were stained with DAPI (blue). The scale bar is 10 µm. The white and open red arrows point to globular VLSs and filamentous structures, respectively. The discontinued yellow line labels the nucleus position. **b)** Immunofluorescence images of LMH cells expressing V5-NSP5 (RVD, RVF) or NSP5-V5 (RVG) (first column) or NSP2-HA (second column) individually, or combined (third-fifth column). At 16 hpt, the cells were fixed and immunostained for detection of NSP5 (anti-V5, green) and NSP2 (anti-HA, red). A merged image is shown in the right column. The scale bar is 10 µm. The white arrows point to VLSs, and the yellow arrows point to globular inclusion formed by NSP5-V5/G.

### Role of NSP5 ordered region across RV species A to J

The NSP5 of RVA has an ordered region at its C-terminus corresponding to an alpha-helix (43, 66). It has been demonstrated that this region, ranging from amino acids 180 to 198 for simian strain SA11, termed tail, is necessary for NSP5 self-oligomerization (34, 45) and association with other RV proteins like NSP2, NSP6, VP2, and VP1 (34, 38, 40, 45, 67). A recombinant RV harboring NSP5 with a deleted tail region has an impaired replication because it cannot form viroplasms (37). Accordingly, we investigated if the ordered region of NSP5 in other RV species plays a role in VLS formation. For this purpose, we identified the ordered region in NSP5 from RV species A to J based on PONDR scores (**Fig 1b-e**) and designed NSP5 deletion mutants lacking their tail region (ΔT) (**Fig 3a and b**). For simplicity, the ordered regions of NSP5/D and F, even if at the N-terminal region, are likewise denominated ΔT. The expression of the recombinant proteins was confirmed by immunoblotting (**Fig 3b**). As expected (15, 32), the expression in MA/cytBirA cells of RVA NSP5ΔT-BAP with NSP2-HA did not allow VLS formation (**Fig 3c**). Similarly, the formation of globular VLS was impaired with NSP5ΔT of the mammalian RV species B and I. The cytosolic distribution of NSP5ΔT-BAP of RV species C, G, H, and J did not change in co-expression with their respective NSP2-HA, similar to the corresponding full-length NSP5-BAP. Avian BAP-ΔTNSP5/D formed VLS, while BAP-ΔTNSP5/F formed distinct nuclear inclusions in MA/cytBirA cells that did not co-localize with NSP2-HA. Independently of the RV species tested, NSP2-HA remained homogenously dispersed in the cytosol when in the presence of NSP5 with the deleted ordered region. A similar pattern of distribution of BAP-ΔTNSP5 of RVD and RVF was observed for LMH cells (**Fig 3d**). Thus, BAP-ΔTNSP5/D forms VLSs, and BAP-ΔTNSP5/F forms both cytosolic and nuclear inclusions when co-expressed with their respective NSP2-HA. NSP5ΔT-V5/G expressed alone in LMH cells is homogenously distributed in the cytosol, contrasting with full-length NSP5-V5/G, which forms cytosolic inclusions (**Fig 3e**). Moreover, the co-expression of NSP5ΔT-V5 and NSP2-HA of RVG had an impaired ability to form VLSs in LMH cells.

**Figure 3.**
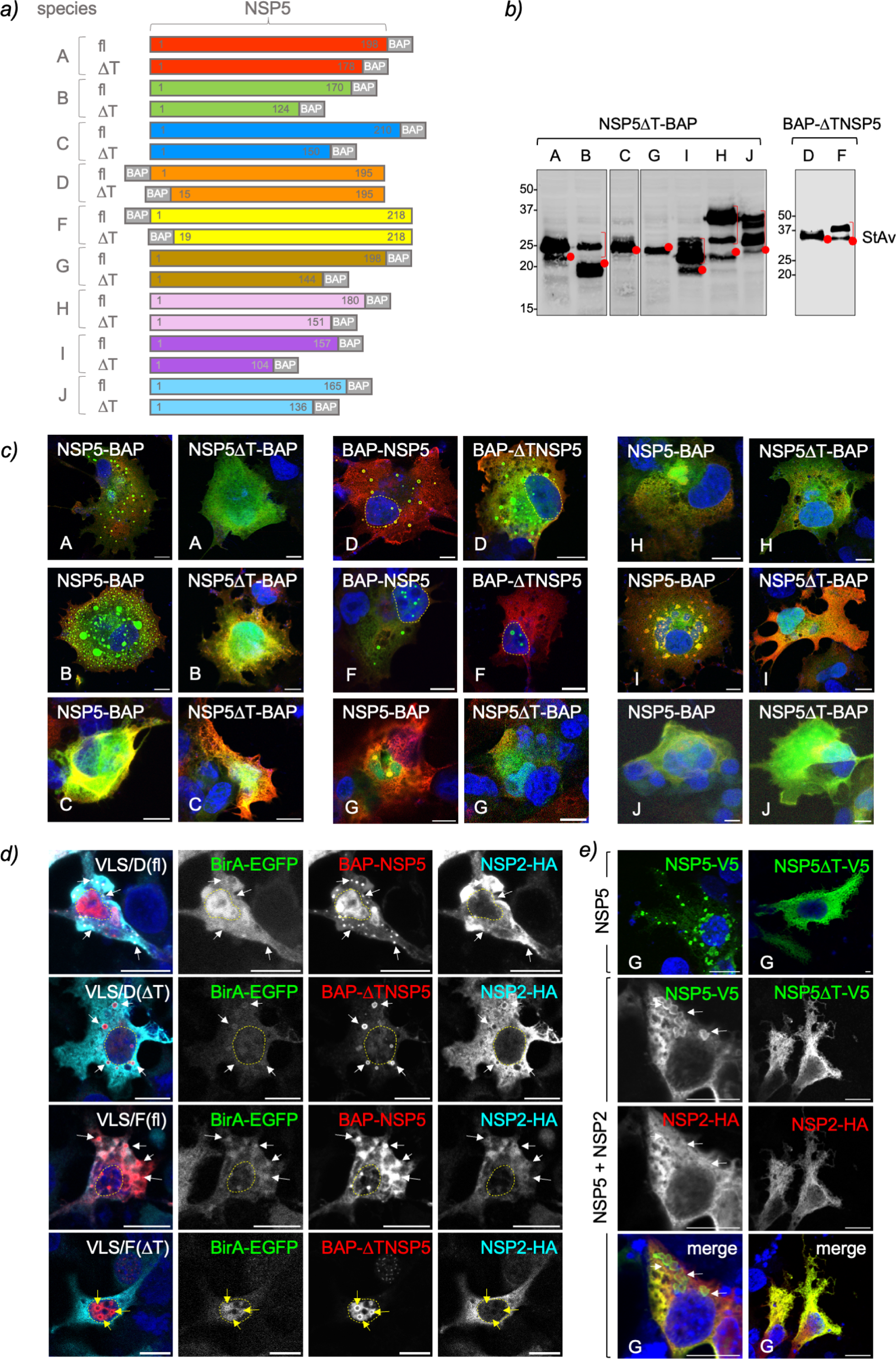
NSP5 ordered region, tail, is required for VLS formation among RV species. **a)** Schematic representation of RV species A to J of full-length (fl) NSP5 and NSP5 with deleted tail region (ΔT) fused to a BAP tag at the N- or C-terminus as indicated. **b)** Immunoblotting of MA/cytBirA cells expressing NSP5ΔT-BAP (RVA to RVC and RVG to RVJ) and BAP-ΔTNSP5 (RVD and RVF). The membrane was incubated with streptavidin-IRDye800. The red dots indicate the predicted molecular weight of the proteins. The red bracket shows a slow migration pattern of the protein. **c)** Immunofluorescence images of MA/cytBirA cells co-expressing NSP2-HA with NSP5 or NSP5ΔT fused to BAP tag. At 16 hpt, the cells were fixed and immunostained for the detection of NSP5 or NSP5ΔT fused to BAP tag (StAv, green) or NSP2-HA (anti-HA, red). Nuclei were stained with DAPI (blue). The scale bar is 10 µm. **d)** Immunofluorescence images of LMH cells co-expressing BirA-EGFP, NSP2-HA with BAP-NSP5 or BAP-ΔTNSP5 of RVD and RVF. The cells were fixed at 16 hpt and immunostained for detection of NSP5 (StAv, red), NSP2 (anti-HA, cyan), and BirA-EGFP (green). Nuclei were stained with DAPI (blue). The scale bar is 10 µm. **e)** Immunofluorescence of LMH cells expressing NSP5-V5/G fl (NSP5-V5) or ΔT (NSP5ΔT-V5) alone or together with NSP2-HA/G. After fixation, the cells were immunostained for the detection of NSP5 (anti-V5, green) and NSP2 (anti-HA, red). Nuclei were stained with DAPI (blue). The scale bar is 10 µm. A merged image for the co-expression of NSP5-V5 with NSP2-HA is shown. In **d**, and **e**, the white arrows point to VLSs, the yellow arrows point to nuclear inclusions, and discontinued yellow lines label the nucleus position.

### NSP5 self-oligomerizes in RV species A to J

A biophysical feature of RVA NSP5 corresponds to its ability to self-oligomerize(34, 45). Additionally, the C-terminal tail of NSP5 seems to be a requirement for VLS formation (37, 45). Therefore, we interrogated whether the tail region of NSP5 in the other RV species is also necessary for self-oligomerization. For this purpose, we added a V5 tag to the full-length NSP5 of all RV species (**Fig S6 a and b**) and co-expressed them with their respective BAP-tagged full-length NSP5 or NSP5ΔT. Next, biotinylated cell extracts were immunoprecipitated with a monoclonal anti-V5 antibody to detect the association between NSP5-V5 with NSP5-BAP or NSP5ΔT-BAP (**Fig 4**). We found that NSP5 of all RV species tested could self-oligomerize, as denoted by the ability of NSP5-V5 to pull down full-length NSP5-BAP. However, NSP5-V5 was unable to pull down NSP5-BAP from RV species having deleted the tail region at the C-terminus (NSP5ΔT-BAP), corresponding to RV species A, B, C, G, H, I, and J (**Fig 4a, b, c, f, g, h, and i**). The association of V5-NSP5 with BAP-NSP5ΔT of RV species D and F (**Fig4 d and e**) remained strong, suggesting no role of their N-terminal region for NSP5 self-oligomerization.

**Figure 4.**
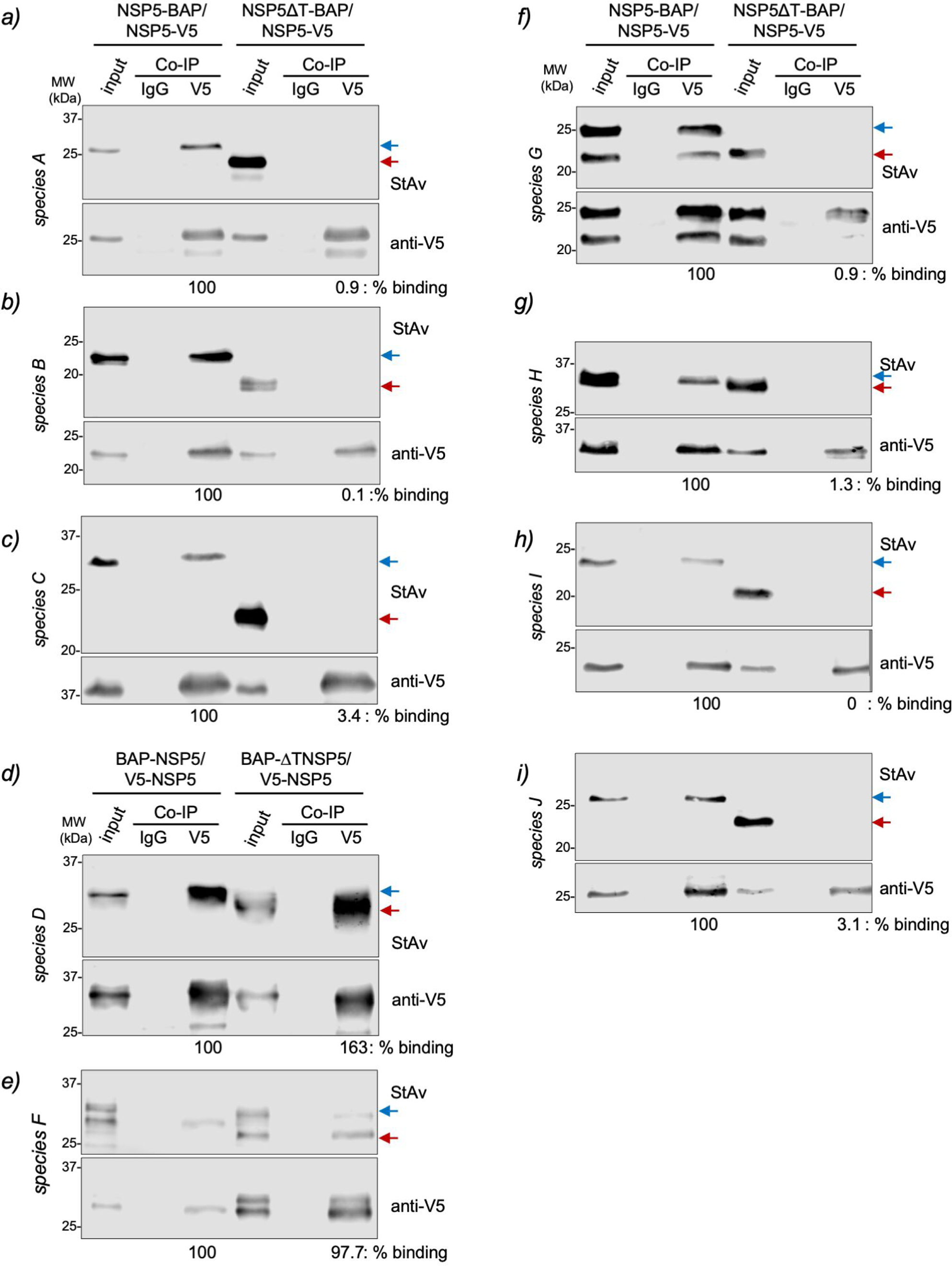
The NSP5 tail of RV species A to C and G to J is necessary for its self-oligomerization.Anti-V5 immunoprecipitated from extracts of MA/cytBirA cells co-expressing full-length NSP5-V5 with full-length NSP5-BAP or NSP5ΔT-BAP for RV species A **(a)**, B **(b)**, C **(c)**, G **(f)**, H **(g)**, I **(h)** and J **(i)**. For RV species D **(d)** and F **(e)**, the cells co-expressed full-length V5-NSP5 with full-length BAP-NSP5 or BAP-ΔTNSP5. The membranes were incubated with streptavidin conjugated to IRDye800 (top panel) and mouse mAb anti-V5 (bottom panel). The input corresponds to 5% of crude cell extract. IgG corresponds to immunoprecipitation with isotype control antibody. Blue and red arrows point to full-length NSP5-BAP and NSP5ΔT-BAP, respectively. The percentage of binding corresponds to the percentage of BAP-tagged NSP5 associated with V5-tagged NSP5.

### NSP2 association with NSP5 across RV species A to J

In RV species A, NSP2 associates with NSP5 by binding at the N-terminal region (amino acids 1 to 33) and its C-terminal tail (amino acids 180-198)(15, 67). In this context, we interrogated whether NSP2 binds to NSP5 in RV species B to J through their NSP5 tail region. For this purpose (**Fig 5**), we co-expressed NSP2-HA with NSP5-BAP or NSP5ΔT-BAP. The biotinylated cell extracts were immunoprecipitated with a monoclonal anti-HA antibody, followed by immunoblotting analysis of the pull-down complexes to detect BAP-tagged full-length NSP5 or NSP5ΔT. As expected (**Fig 5a**), NSP2 RVA binds to full-length NSP5-BAP but also to NSP5ΔT-BAP (15). Except for NSP2/J (**Fig 5i**), NSP2 of all RV species tested bind their full-length NSP5. Interestingly (**Fig 5b, c, d, e, and f**), NSP2-HA of RV species B, C, D, F, and G had a drastic impairment of the binding to BAP-tagged NSP5ΔT, suggesting that NSP2 binds specifically to the predicted ordered regions of NSP5. Similar to NSP2-HA/A (**Fig 5g and h**), the NSP2-HA of RV species H and I do not show a reduction in the binding to NSP5 (depicted in **Fig 5** as the ratio of NSP5/NSP2 binding) when co-expressed with their respective NSP5ΔT-BAP, suggesting that in these proteins, NSP2 binds to an additional region.

**Figure 5.**
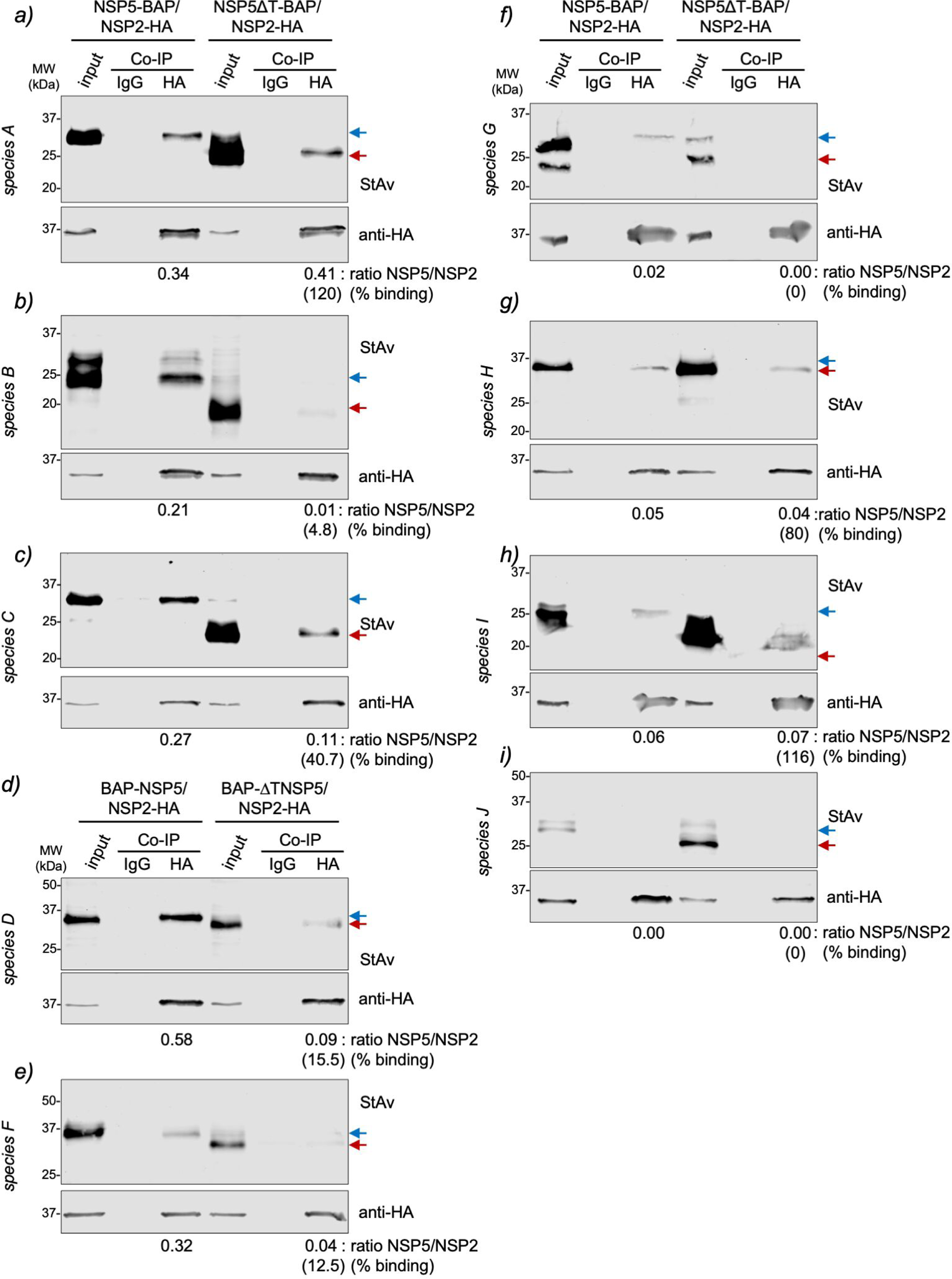
NSP5 requirements for association with NSP2 across RV species A to J. Anti-HA immunoprecipitated from extracts of MA/cytBirA cells co-expressing NSP2-HA with full-length NSP5-BAP or NSP5ΔT-BAP for RV species A **(a)**, B **(b)**, C **(c)**, G **(f)**, H **(g)**, I **(h)**, and J **(i)**. For RV species D **(d)** and F **(e)**, the cells co-expressed NSP2-HA with full-length BAP-NSP5 or BAP-NSP5ΔT. The membranes were incubated with streptavidin conjugated to IRDye800 (top panel) and mouse mAb anti-HA (bottom panel). The input corresponds to 5% of crude cell extract. IgG corresponds to immunoprecipitation with isotype control antibody. Blue and red arrows point to full-length NSP5-BAP and NSP5ΔT-BAP, respectively. The NSP5/NSP2 association ratio of BAP-tagged full-length NSP5 and NSP5ΔT to NSP2-HA is indicated. The percentage of binding of NSP5ΔT to NSP2 relative to full-length NSP5 binding to NSP2 is indicated.

### Heterologous formation of VLS between RV species

Since some common elements are present among the diverse RV species, such as the ordered regions, the ability of NSP5 to oligomerize, and the capacity of some NSP2 species to form octameric structures, we assessed if these common elements allow interspecies VLS formation. In the first instance (**Fig S7a and Table 3**), we co-expressed NSP5/A with NSP2-HA of RV species A to J and checked for the formation of VLSs. As expected, homologous formation of VLS is observed for co-expression of NSP5 and NSP2-HA of RVA. Interestingly, NSP5/A forms globular VLS with NSP2-HA/I and filamentous and globular-like structures with NSP2-HA/C. Indeed, the combinations of NSP5/A with NSP2-HA of RV species B, D, F, G, H, and J did not lead to VLS formation, and the proteins remained homogenously distributed in the cytosol of the cell. Next (**Fig S7b and Table 3**), we expressed NSP5-BAP of the RV species A to J with NSP2-HA/A. Only homologous pairs of NSP5-BAP with NSP2-HA of RVA formed VLSs in these specific conditions. Instead, the NSP5-BAP of the other RV species behaved as expressed without NSP2-HA (**Fig 2a**).

**Table 3.** Formation of heterologous RV VLS.

| co-expression<sup>a</sup> |  | VLS<br>formation<sup>c</sup> | co-expression<sup>b</sup> |  | VLS<br>formation |
| --- | --- | --- | --- | --- | --- |
| NSP5<br>species | NSP2-HA<br>species |  | NSP5-BAP<br>species | NSP2-HA<br>species |  |
| A | A | + | A | A | + |
| A | B | - | B | A | - |
| A | C | filamentous | C | A | - |
| A | D | — | D | A | -, nuclear<br>aggregates |
| A | F | - | F | A | -, nuclear<br>inclusions |
| A | G | - | G | A | -, aggregates |
| A | H | - | H | A | - |
| A | I | + | I | A | - |
| A | J | - | J | A | - |

- 929 a) Confirmed by co-immunostaining of NSP5 of RVA and NSP2 HA of species A-J using specific guinea pig pAb anti-NSP5 and mAb anti-HA, respectively. - 931 b) Confirmed by co-immunostaining of NSP5-BAP of species A to J and NSP2-HA of RV species A using StAv and mAb anti-HA, respectively. c) The result corresponds to the observation of three independent experiments.

Next, we compared the evolutionary proximity of NSP5 and NSP2 of the diverse RV species (**Fig 6**). Interestingly, the phylogenetic trees of NSP5 and NSP2 present similarities regarding the common ancestor for the analyzed RV species. Like that, it is possible to form couples of RV species based on their closer common ancestors. Thus, RVA is closer to RVC, RVB is closer to RVG, RVD is closer to RVF, and RVH is closer to RVJ. Meanwhile, RVI shared a common ancestor with RVB, RVG, RVJ, and RVH. In this context (**Fig 7a**), we assessed VLS formation by co-expressing NSP5-BAP and NSP2-HA of RV species A and C in all four interspecies combinations of NSP2 and NSP5 (A/A, C/C, A/C, and C/A). Only NSP5-BAP/A with NSP2-HA/A formed globular cytosolic VLS, while the other combinations formed filamentous VLSs. The combinations of NSP5-BAP and NSP2-HA of RVB and RVG (**Fig 7b**) showed VLS formation by co-expression of NSP5-BAP/B or NSP5-BAP/G with NSP2-HA/G. Interestingly (**Fig 7c**), avian BAP-NSP5 and NSP2-HA of RVD and RVF formed VLS in all the combinations. As expected (**Fig 7d**), the combinations of NSP5-BAP with NSP2-HA of RV species H and J did not lead to globular VLS formation.

**Figure 6.**
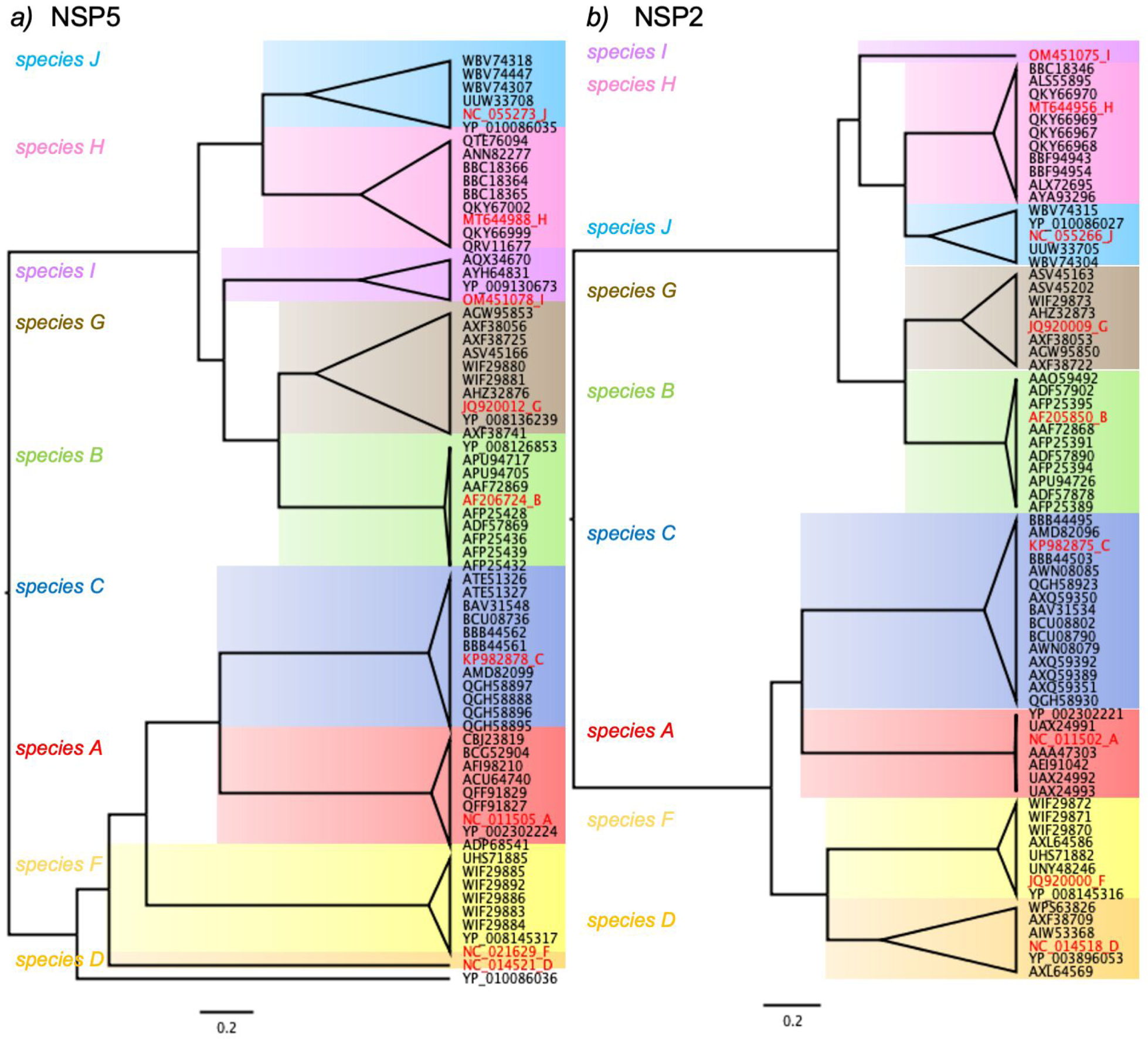
Phylogenetic trees for amino acid sequences of NSP5 and NSP2 in RV species A to J. Maximum likelihood tree showing phylogenetic relationships between NSP5 **(a)** and NSP2 **(b)** genes of RV species A to J. The red label corresponds to the coding sequences used in this study, followed by a letter associated with its RV species. Each RV species has a colored panel: RVA, red; RVB, green; RVC, blue; RVD, orange; RVF, yellow; RVG, brown; RVH, pink; RVI, violet; and RVJ, light blue. The scale is 0.2 substitutions per nucleotide.

**Figure 7.**
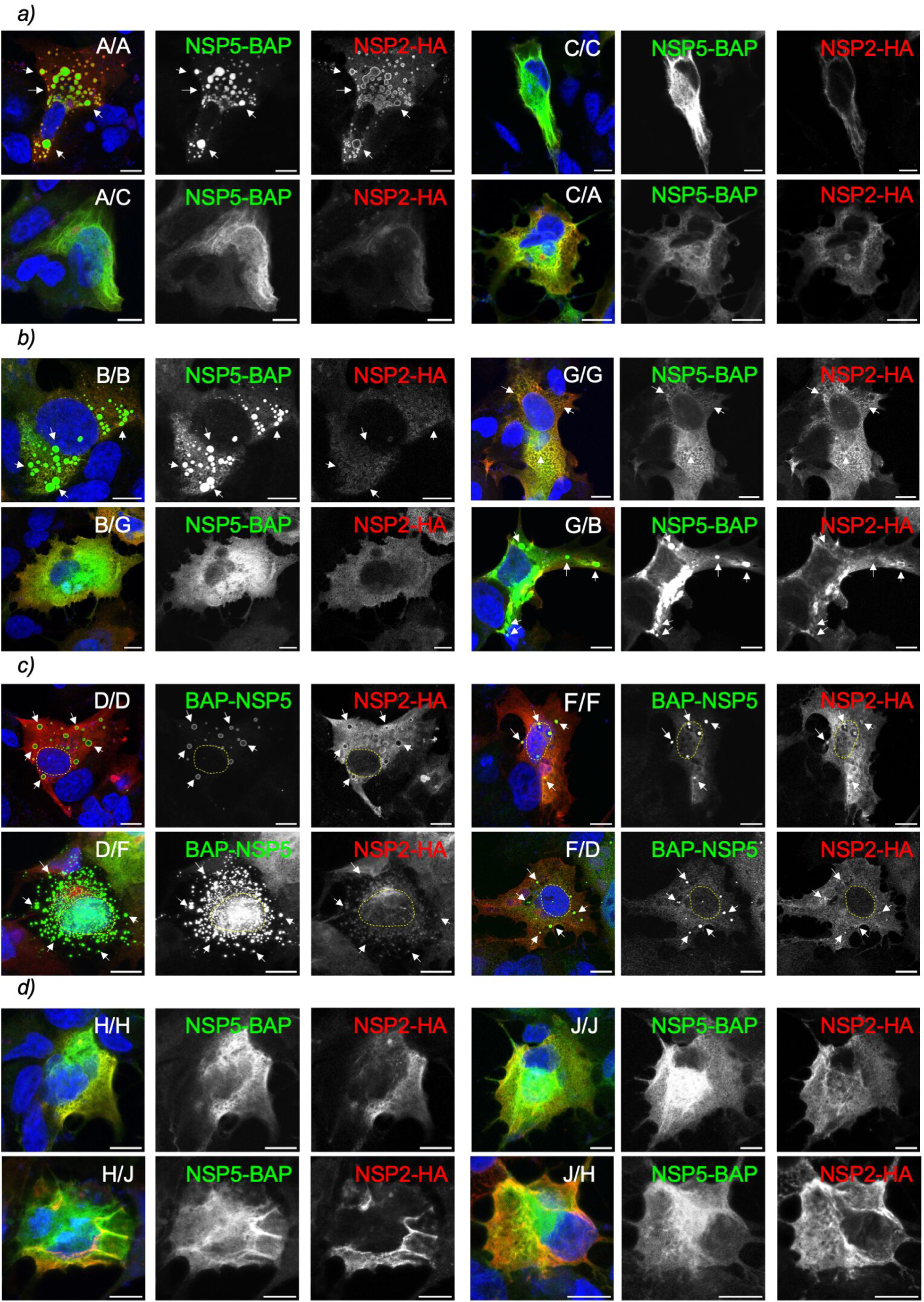
Heterologous formation of VLS among closely related RV species. Immunofluorescence images of MA/cytBirA cells co-expressing NSP5 tagged to BAP and NSP2-HA of closely related RV species A and C **(a)**, B and G **(b)**, D and F **(c)**, and H and J **(d)**. After fixation, the cells were immunostained for the detection of NSP5 (StAv, green) and NSP2 (anti-HA, red). The nuclei were stained with DAPI (blue). The indication at the top right corner corresponds to the RV species of NSP5 and NSP2, respectively. The scale bar is 10 µm. The white arrows point to globular VLSs. The yellow lines label the nucleus position.

### Chimeric NSP5ΔT/H and J with tail of NSP5/A form VLS

Our results demonstrated that NSP5/H and J were unable to form VLS with their respective NSP2, even though their tail regions appear to be necessary for oligomerization (Fig 4g and i). However, it seems that the tail regions alone are not sufficient for a strong interaction with NSP2 (**Fig 5 g and i**). Additionally, we have shown that NSP5 and NSP2 from certain RV species can be swapped (**Fig 7**), which allows for VLS formation. In addition, it has been shown that the short tail region of NSP5/A (amino acids 180-198) is essential for viroplasm formation (37). We next investigated whether replacing the predicted ordered region of NSP5/H and NSP5/J with the tail of NSP5 (TA) would enable the VLS formation with NSP2 of their respective species. For this purpose (**Fig 8a and Fig S5 c and d**), we built two chimeric proteins harboring NSP5ΔT/H or NSP5ΔT/J fused to the TA followed by a BAP-tag, termed NSP5ΔT/H/TA-BAP and NSP5ΔT/J/TA-BAP, respectively. Additionally, we built a third chimeric protein composed of full-length NSP5/H fused to TA (NSP5/H/TA-BAP). Interestingly (**Fig 8b**), NSP5ΔT/H/TA-BAP and NSP5/H/TA-BAP expressed alone were found to be homogeneously dispersed in the cytosol of the cells, whereas NSP5ΔT/J/TA formed discrete cytosolic inclusion resembling VLSs. As expected, VLSs are not formed by the co-expression of NSP5-BAP/H with either NSP2-HA/A or NSP2-HA/H (**Fig 8c, top panel**), the co-expression of NSP5-BAP/J with either NSP2-HA/A or NSP2-HA/J (**Fig 8d, top panel**), or the co-expression of NSP2-HA/A with either NSP5ΔT/H/TA-BAP or NSP5ΔT/J/TA-BAP (**Fig 8c and d, top-middle panels**). Indeed, the co-expression of chimeric NSP5ΔT/H/TA-BAP with NSP2-HA/H (**Fig 8c, middle panel**) or NSP5ΔT/J/TA-BAP with NSP2-HA/J (**Fig 8d, bottom panel**) resulted in well-discernible globular VLSs. Moreover, adding TA to the full-length NSP5/H when co-expressed with NSP2-HA/H also allowed the formation of VLSs (**Fig 8c, bottom panel**). Using an immunoprecipitation assay, we then tested the ability of these chimeric proteins to associate with their respective NSP2s. Our results demonstrated that NSP2-HA/H binds to BAP-tagged NSP5/H, NSP5ΔT/H/TA, and NSP5/H/TA (**Fig 8e and f**). However, this binding was apparently weaker for NSP5/H/TA-BAP and NSP5ΔT/H/TA-BAP than for NSP5-BAP/H. Interestingly (**Fig 8g**), the chimeric NSP5ΔT/J/TA-BAP can associate with NSP2-HA/J, contrasting the wild-type NSP5-BAP/J, which cannot bind to NSP2-HA/J.

**Figure 8.**
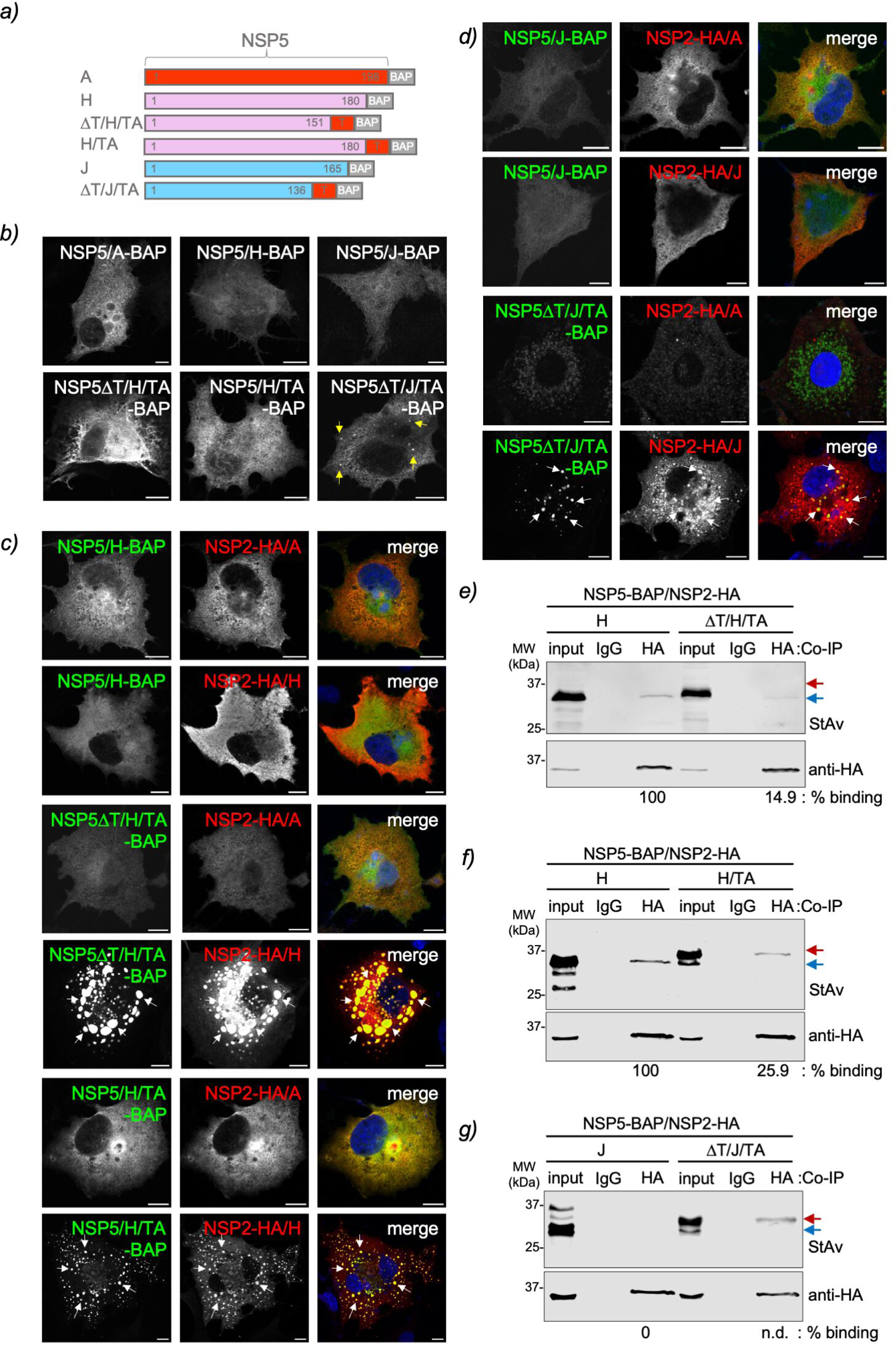
Chimeric NSP5 RVH and RVJ harboring RVA tail form VLS. **a)** Schematic representation of chimeric NSP5 RVH and RVJ with the NSP5 RVA (TA) tail fused to a BAP tag at the C-terminal region. **b)** Immunofluorescence images of MA/cytBirA cells expressing wild-type NSP5 and chimeric NSP5 of RVH and RVJ. The yellow arrows point to globular cytosolic inclusions. **c)** Immunofluorescence images of MA/cytBirA cells co-expressing NSP5/H-BAP (top panel), NSP5ΔT/H/TA-BAP (middle panel) and NSP5/H/TA (bottom panel) with either NSP2-HA/A or NSP2-HA/H. **d)** Immunofluorescence images of MA/cytBirA cells co-expressing NSP5/J-BAP (top panel) and NSP5ΔT/J/TA-BAP (bottom panel) with either NSP2-HA/A or NSP2-HA/J. The cells were fixed at 16 hpt and immunostained for the detection of NSP5 (StAv, green) and NSP2 (anti-HA, red). The nuclei were stained with DAPI (blue). The scale bar is 10 µm. The white arrows point to globular VLSs. Anti-HA immunoprecipitated from extracts of MA/cytBirA cell co-expressing NSP2-HA/H with NSP5ΔT/H/TA **(e) or** NSP5-H/TA **(f)** and co-expressing NSP2-HA/J with NSP5ΔT/J/TA **(g)**. As a control, wild-type NSP5 for each tested RV species is included. The membranes were incubated with streptavidin conjugated to IRDye800 (top panel) and mouse mAb anti-HA (bottom panel). The input corresponds to 5% of crude cell extract. IgG corresponds to immunoprecipitation with isotype control antibody. Blue and red arrows point to full-length NSP5-BAP and NSP5ΔT-BAP, respectively. The percentage of binding of BAP-tagged full-length NSP5 and NSP5ΔT to NSP2-HA are indicated.

## Discussion

A recurrent question in the rotavirus research field relates to the formation of viroplasms in non-RVA species. The accepted ICTV RV species A to J share common features, such as eleven dsRNA genome segments with conserved 5’ and 3’-untranslated sequences. Moreover, the open reading frames on these genome segments encode proteins with primary amino acid sequences homologous to RVA species, including the proteins involved in viroplasm formation (NSP5, NSP2, and VP2) and replication (VP1, VP2, VP3, VP6). Even if all the elements that facilitate the formation of viroplasms are present, it is still cumbersome to demonstrate their existence because of the lack of tools for viroplasm recognition, like specific antibodies targeting proteins of every RV species. Additionally, these RV species are not replicating in tissue culture, making it challenging to identify viroplasms using electron microscopy techniques. In this study, we extrapolated the knowledge acquired from RVA viroplasms and used the co-expression of NSP5 and NSP2 of the RV species B to J to detect the formation of globular VLSs. These structures, even if unable to replicate, have been demonstrated to be simplified models for studying highly complex viroplasms in RVA (15, 16, 31, 33, 40, 51, 68), allowing even the investigation of the recruitment of cellular proteins as was recently demonstrated with the TRiC chaperonin (65). As summarized in **Table 4**, our experiments revealed that NSP5 and NSP2 of species B, D, F, G, and I form VLSs that are morphologically similar to those observed for RVA. We found that only some of the RV species, specifically C, H, and J, were unable to form globular VLS upon co-expression of NSP5 with NSP2. For NSP5 and NSP2 of RVC, we observed a mixed population of structures, globular and filamentous, that depended on the expression levels of the two proteins, with a shift towards globular structures when overexpressing NSP2. In the standard conditions for detecting VLS formation, the RVC pair formed nocodazole-sensitive filamentous structures, suggesting direct dependence on the MT network. It is not surprising to observe filamentous VLS since viral factories of some members of the *Reoviridae* family also form filamentous structures. For example, the µ2 protein of mammalian orthoreovirus (MRV) strain T1L, which can bind MTs, forms filamentous viral factories, while the µ2 protein of strain T3D^N^, which cannot bind to MTs, forms globular viral factories (69). Additionally, we found that VLS formation in avian RV species D, F, and G is host-dependent since these structures formed better in avian LMH cells than in mammalian MA104 cells. It is particularly true for RVG, where NSP5 expressed alone forms aggregates in MA104 cells, while in LMH cells, it forms regularly shaped globular inclusions. Also, NSP5 of RVF is an interesting case since it forms globular nuclear inclusions when expressed alone in MA104 cells but not in LMH cells. The ability of NSP5 RVF and RVG to form inclusions per se is a novelty in the RV field, however, not for the *Reoviridae* family where, for example, the MRV protein µNS forms cytosolic inclusions when expressed in the absence of other virus proteins (70, 71). The NSP5-NSP2 couples of RVH and RVJ do not lead to VLS formation at any of the tested ratios, indicating that either these two proteins are not sufficient to form VLS and require additional RV proteins like VP2 (31, 40) or that viroplasms in these species are composed of other RV proteins not related to NSP5 or NSP2.

**Table 4.** Summary of NSP5 requirements across RV species A to J.

| RV species | globular VLS formation<sup>a</sup> |  | NSP5 oligomerization<sup>b</sup> |  | NSP5 interaction with NSP2<sup>c</sup> |  |
| --- | --- | --- | --- | --- | --- | --- |
|  | fl NSP5<sup>g</sup> | NSP5ΔT | fl NSP5 | NSP5ΔT | fl NSP5 | NSP5ΔT |
| **A** | + | - | + | - | + | + |
| **B** | + | - | + | - | + | +/- |
| **C** | -<sup>d</sup> | - | + | - | + | + |
| **D** | + | + | + | + | + | +/- |
| **F** | + | +<sup>e</sup> | + | + | + | +/- |
| **G** | +<sup>f</sup> | - | + | - | +/- | - |
| **H** | - | - | + | - | +/- | +/- |
| **I** | + | - | + | - | +/- | +/- |
| **J** | - | - | + | - | - | - |

a) Confirmed by co-immunostaining for detection of both VLS components, NSP5 and NSP2
(transfection ratio 1:2=NSP5:NSP2).
b) Confirmed by co-immunoprecipitation of NSP5-V5 using anti-V5 antibody.
c) Confirmed by co-immunoprecipitation of NSP2-HA using anti-HA antibody. Weak association determined by an NSP5/NSP2 ratio below 0.1.
d) Form filamentous cytosolic structures. Upon overexpression of NSP2 (ratio 1:8=NSP5:NSP2), globular VLS can be detected.
e) Forms nuclear inclusions.
f) Forms VLS only in chicken epithelial LMH cells.
g) (+) positive, (+/-) weak and (-) negative.

Interestingly, the NSP5 proteins of all the RV species tested in this study were predicted to be intrinsically disordered, similar to NSP5 of RV species A. In this context, it is known that NSP5/A can self-oligomerize, a feature that is conserved in NSP5 of species A to J, as demonstrated by co-immunoprecipitation assays. The self-oligomerization of NSP5 in RVA through its tail is essential for viroplasm formation (15, 37). Thus, recombinant RVA with the tail of NSP5 deleted does neither form viroplasms nor replicate if not rescued by a cell line expressing full-length NSP5 (37). Additionally, the tail of NSP5 is required to associate with other virus proteins such as NSP2 (15), VP1 (38), VP2 (40) or NSP6 (34). In this context, the deletion of the tail in NSP5 hampered self-oligomerization in those RV species that have an ordered region at the C-terminal end (RV species A, B, G, and I), which is directly correlated with their ability to form globular VLSs. However, deleting the predicted ordered NSP5 N-terminal region of RV species D and F did not affect oligomerization and the ability to form globular inclusions. Interestingly, the deletion of the N-terminus of NSP5/F led to the formation of nuclear inclusions even in co-expression with NSP2. This result suggests that the binding site to NSP2 is localized at its N-terminal region, that NSP2 retains NSP5/F in the cytosol, and that the ability to form inclusions depends on the center or C-terminal region of NSP5/F. Consistent with our results, AlphaFold3 predicts **(Fig. S7)** that the full-length NSP5 dimerizes through its intertwined C-terminal tail in RV species A, B, C, G, H, I, and J. The deletion of this tail appears to disrupt the intertwined interaction between the NSP5 monomeric structures, hampering their dimerization. However, the intertwined tail between NSP5/D and F is not observed by AlphaFold3 at either N- or C-terminal regions, suggesting that these proteins oligomerize through another undetermined region.

In RVA, NSP2 associates with NSP5 through its N-terminus (amino acid region 1-33) and tail (amino acid region 190-198)(15, 67). The biochemical properties of NSP5 and NSP2, requiring transfer of nucleotides and high peptide chain flexibility (43, 50), are consistent with the fact that their association needs to be stabilized by chemical crosslinking to support detection. Similarly, the NSP2 and NSP5 association must be chemically stabilized in RV species B to I for detection. Even though the association of NSP5 to NSP2 seems eight times stronger for RV species A, B, C, D, and F than RV species H, G, and I, as denoted by the estimation of their relative binding ratios. Interestingly, NSP2/J does not associate with NSP5 at any of the tested conditions of the DSP crosslinker. As described previously using crystallography and cryo-electron microscopy, NSP2 of RV species A (49), B (61), and C (62) has octameric structures with a 4-2-2 disposition. Consistent with these observations, AlphaFold3 also predicted that NSP2 of RV species A to J is organized as octamers with a 4-2-2 crystal symmetry **(Fig S3)**. These outcomes suggest that NSP5 is responsible for orchestrating the association with NSP2. It is noticed that a positive association of NSP5 and NSP2 does not necessarily lead to globular VLS formation, for example, for the couple of RVC (**Table 4**). As expected, the deletion of the NSP5 tail does not disrupt the association with NSP2, for example, for RVA, RVC, RVH, and RVI, probably due to association with another region of NSP5. However, the association was drastically hampered in RV species D, F, and G when the ordered region of NSP5 was deleted.

An interesting aspect relates to the interspecies formation of NSP5 and NSP2 VLSs. Only specific close-related RV species can interchange these proteins to form globular VLS. The RV species B and G, along with the avian RV species D and F, have been shown to possess promiscuous NSP5 and NSP2 proteins. As a result, they can form globular VLS with any combination of NSP5 and NSP2. This result suggests that at least in avian RVs, interspecies reassortment is possible. The use of chimeric NSP5/H and J, having incorporated the tail of NSP5/A, allowed the formation of VLS exclusively with their cognate NSP2 but not with NSP2-HA/A. This indicates that cognate couples of NSP2 and NSP5 associate specifically in a region independent of their NSP5 tail. Moreover, the NSP5 tail A stabilizes these associations, suggesting that other RV proteins could favor RV species H and J in forming viroplasms. It is well known that VP2 also allows the formation of VLS (31, 40). Nevertheless, VP6 also improves the stabilization of viroplasms, at least in RV species A (68). In this context, we cannot exclude that post-translational modifications, such as specific phosphorylation of both NSP5 and NSP2, could also influence the formation of VLS and viroplasms in non-RVA species. The non-canonical site for casein kinase 1-alpha (**Fig S1a**) that is primed in NSP5 serine 67 of RVA strain SA11 is conserved in RV species A, C, D, and F. Surprisingly, the closely related RV species B and G have replaced the serine 67 by a tyrosine, which may be phosphorylated by CK1-alpha. RV species H, I, and J have obliterated this phosphorylation site. Coincidentally, the NSP2 phosphorylation site for CK1-alpha in RVA corresponding to serine 313 is also conserved in RV species A, C, D, and F (**Fig S1b**). Additional studies need to be performed to prove the role of these phosphorylation sites in VLS and viroplasm formation across the RV species.

## Materials and Methods

### Cells and viruses

MA104 cells (embryonic rhesus monkey kidney, ATCC^®^CRL-2378, RRID: CVCL_3845) were cultured in Dulbecco’s modified Eagle’s medium (DMEM, Gibco^®^BRL) supplemented with 10% fetal calf serum (FCS)(AMIMED, BioConcept, Switzerland) and penicillin (100 U/ml)-streptomycin (100 µg/ml)(Gibco, Life Technologies). MA/cytBirA (24) cell lines were grown in DMEM supplemented with 10% FCS, penicillin (100 U/ml)-streptomycin (100µg/ml), and 5µg/ml puromycin (InvivoGen, France). LMH cells (chicken hepatocellular carcinoma epithelial, ATCC^®^CRL2117) were cultured in Waymouth’s MB752/1 (Sartorius) medium supplemented with 10% FCS and penicillin (100 U/ml)-streptomycin (100 µg/ml).

The recombinant vaccinia virus encoding T_7_ RNA polymerase (strain vvT7.3) was amplified as previously described (72).

### Antibodies and reagents

Guinea pig anti-NSP5, mouse monoclonal (mAb) anti-NSP5 (clone 2D2), and guinea pig anti-NSP2 were described previously (15, 64). Mouse mAb anti-HA (clone HA-7) and mouse anti-tubulin (clone B5-1-12) were purchased from Merck. Mouse mAb anti-V5 (SV5-PK1) was purchased from Abcam. Mouse MAb anti-α-tubulin was directly conjugated to Atto 488 using the Lightning-Link Atto 488 conjugation kit from Innova Bioscience, United Kingdom. Alexa Fluor594 anti-HA.11 (clone 16B12) was purchased from BioLegend. Streptavidin-DyLight™ 488 and mouse secondary antibodies conjugated to Alexa 488 or Alexa 594 were purchased from Thermo Fisher Scientific. Streptavidin-IRDye®800CW and secondary antibodies for immunoblot conjugated to IRDye^®^680 and IRDye^®^800CW were purchased from LI-COR.

Nocodazole was purchased from Sigma. Dithio(succinimidylpropionate)) was purchased from ThermoFisher Scientific.

### Rotavirus sequences

The rotavirus NSP5 and NSP2 open reading frames from species B to J were synthesized at GenScript using the sequences described in the supplemental material. NSP5 B to J were cloned in pCI-Neo (Promega) between *Mlu*I/*Not*I restriction sites. NSP2 B to J were cloned in pCI-Neo (Promega) between *EcoR*I/*Not*I restriction sites. The following sequences for NSP5 and NSP2 were used in this study (GenBank accession number listed for each): RVB NSP5 human strain CAL-1(AFE206724), RVB NSP2 human strain CAL-1(AF205850), RVC NSP5 porcine strain 12R021 (KP982878), RVC NSP2 porcine strain 12R021(KP982875), RVD NSP5 chicken strain 05V0049(NC_014521), RVD NSP2 chicken strain 05V0049(NC_014518), RVF NSP5 chicken strain 03V0568(NC_021629), RVF NSP2 chicken strain 03V0568(JQ920000), RVG NSP5 chicken strain 03V0567(JQ920012), RVG NSP2 chicken strain 03V0567(JQ920009), RVH NSP5 porcine strain SP-VC36 (MT644988), RVH NSP2 porcine strain SP-VC36(MT644956), RVI NSP5 raccoon dog strain SD-MO2(OM451078), RVI NSP2 raccoon dog strain SD-MO2(OM451075), RVJ NSP5 bat strain BO4351(NC_055273) and RVJ NSP2 bat strain BO4351(NC_055266).

### Plasmid constructs

pCI-NSP5-BAP/A, B, C, G, H, I, and J were obtained by PCR amplification of pCI-NSP5/A, B, C, G, H, I, and J using specific primers to insert *Mlu*I and BAP tag/*Not*I sites, followed by ligation into those sites in pCI-Neo (Promega). pCI-NSP5-BAP/D and F were obtained by PCR amplification of pCI-NSP5/D and F using specific primers to insert *Mlu*I/BAP tag and *Not*I sites, followed by ligation into those sites in pCI-Neo. pCI-NSP5(1-178)-BAP/A, pCI-NSP5(1-124)-BAP/B, pCI-NSP5(1-150)-BAP/C, pCI-NSP5(1-144)-BAP/G, pCI-NSP5(1-151)-BAP/H, pCI-NSP5(1-104)-BAP/I and pCI-NSP5(1-136)-BAP/J were obtained by PCR amplification of the respective pCI-NSP5-BAP using specific primers to insert indicated ORF deletions and *Mlu*I/*BspE*I restriction enzyme sites, followed by ligation into those sites in their respective in pCI-NSP5-BAP/A. pCI-BAP-NSP5(15-195)/D and pCI-BAP-NSP5(15-195)/F were obtained by PCR amplification of the respective pCI-BAP-NSP5 using specific primers to insert the BAP tag and *Mlu*I/*Not*I restriction sites, followed by ligation into those sites in pCI-Neo. pCI-NSP5-V5/A, B, C, G, H, I and J pCI-NSP5ΔT-V5/A, B, C, G, H, I and J were obtained by annealing the following oligonucleotides 5’-ccggaggcaagcctattcctaaccctctgctgggcctggacagcacctaagc-3’ and 5’-ggccgcttaggtgctgt ccaggcccagcagagggttaggaataggcttgcct-3’ encoding a V5 tag and ligation between *BspE*I and *Not*I restriction sites of ther respective pCI-NSP5-BAP or pCI-NSP5ΔT. pCI-V5-NSP5/D and F were obtained by PCR amplification of ther respective pCI-NSP5 using specific primers to insert *Mlu*I-V5 tag-*Not*I, followed by ligation between *Mlu*I and *Not*I in pCI-Neo. The chimeric NSP5ΔT/H/TA, NSP5/H/TA, and NSP5ΔT/J/TA were synthesized at Genscript as gene fragments and cloned in pCI-NSP5/A-BAP between *Mlu*I and *BspE*I restriction sites. The synthetic chimeric nucleotide sequences are available in the **Supplementary information**.

pCI-NSP2-HA/A was obtained by PCR amplification of pcDNA-NSP2 (33) using specific primers to insert *Xho*I and HA tag/*Not*I, followed by ligation on those sites in pCI-Neo. pCI-NSP2-HA/B, C, D, F, G, H, I, and J were obtained by PCR amplification of their respective pCI-NSP2 using specific primers to insert *Mlu*I and HAtag/*Not*I, followed by ligation on those sites in pCI-Neo. The oligonucleotides were synthesized at Microsynth (Switzerland) and described in **Table S1**.

### AlphaFold predictions

Protein structures of NSP5 monomers, NSP5 dimers, or NSP2 multimers were predicted using AlphaFold3 Server (https://golgi.sandbox.google.com/about)(63).

### IDR prediction

The intrinsically disordered regions of proteins were determined using PONDR® (Molecular Kinetics, Inc, http://www.pondr.com) using the VSL2 algorithm. Data were plotted with GraphPad Prism (version 10.1.1(270)).

### Immunofluorescence

MA/cytBirA cells were seeded at 1×10^5^ cells per well onto coverslips in a 24-well multiwell plate. The cells were infected with vvT7.3, followed by transfection using Lipofectamine 2000 (Thermo Fisher Scientific) as described previously (40). Specifically, cells were transfected with a ratio of NSP5 and NSP2 of 1:2 using 750 ng and 1500 ng of DNA plasmids, respectively. The cells were supplemented with 100 µM biotin when the transfection mixture was added. LMH cells were seeded at a density of 3×10^5^ cells per well onto coverslips in a 24-well multiwell plate. The cells were transfected using *Trans*IT^®^-2020 transfection reagent (Mirus Bio™) according to manufacturer instructions. Briefly, cells were transfected with a ratio of NSP5 and NSP2 of 1:2 using 750 ng and 1500 ng DNA plasmids, respectively. For this purpose, 100 µl Opti-MEM™ (Thermo Fisher Scientific) were mixed with 3 µl of *Trans*IT^®^-2020 and DNA plasmids and incubated for 15 min at room temperature. The cells were washed once with phosphate-buffered saline (PBS) and then, added 400 µl of serum-free Waymouth’s MB725/1 medium per well, followed by the addition of 100 µl transfection mix.

At 16 hpt, the medium was removed, and the cells were fixed with 2% paraformaldehyde for 10 min at room temperature or with ice-cold methanol for 3 min at −20 °C. The cells were permeabilized with 0.01% Triton X-100-PBS for 5 min at room temperature and blocked in 1%BSA-PBS for 20 min at room temperature. The primary and secondary antibodies were diluted in 1% BSA-PBS and incubated for 40 min at room temperature in a humid chamber. The coverslips were mounted onto slides using ProLong™ Gold antifade mountant (Thermo Fisher Scientific).

Images were acquired using a confocal laser scanning microscope (CLSM) (DM550Q, Leica). Data were analyzed with Leica Application Suite (Mannheim; Germany) and ImageJ2 (version: 2.14.0/1.54f, http://imagej.net/Contributors).

### Co-immunoprecipitation and immunoblotting

1.2×10^6^ MA/cytBirA cells were infected with vvT7.3 at a MOI of 3 PFU/cell. Then, the cells were transfected with Lipofectamine 2000 (Thermo Fisher Scientific) in a ratio 1:1 of NSP5-BAP and NSP5-V5 or 1:2 of NSP5-BAP and NSP2-HA following manufacturer instructions. After adding the transfection mixture, the cells were immediately supplemented with 100 µM biotin. At 16 hpt, the cells were lysed in 180 µl of TNN buffer (100 mM Tris-HCl, pH 8.0, 250 mM NaCl, 0.5% Nonidet P-40 and cOmplete protease inhibitor cocktail [Roche, Switzerland]) for 10 min on ice. For NSP5-BAP and NSP2-HA assays, the cells were crosslinked with 300 µM or 600 µM DSP (Dithio(succinimidyl propionate)) prior to lysis as described in detail by Eichwald et al., 2004 (15). The cell lysates were clarified by centrifugation at 17’000 *x g* for 7 min and 4°C and then transferred to a new 1.5 ml tube. The input corresponded to 15 µl of cell lysate. For immunoprecipitation, the cell lysate was split into equal volumes and combined with 2 µg mouse mAb anti-V5 (SV5-PK1) (Abcam, ab27671), mouse mAb anti-HA (clone HA-7)(Merck, H3663) or 2 µg mouse IgG2a kappa isotype control (clone eBM2a) (eBioscience, 14-4724-82) and incubated at 4°C for 30 min with rotation. The cell lysates were combined with 50 µl of Protein G Dynabeads (ThermoFisher, 10004D), equilibrated in TNN, and re-incubated at 4°C for 30 min with rotation. The bead-bound antibody-antigen complexes were washed four times with 500 µl TNN, eluted with SDS sample buffer, and resolved by SDS-PAGE. The proteins were detected by immunoblotting, as described below.

### Immunoblotting

Cells seeded in 12-well tissue culture plates at a density of 2×10^5^ cells per well were lysed directly by adding 25 µl of Laemmli sample buffer 4X (8 % SDS, 40 % glycerol, 200 mM Tris-HCl pH 6.8, 0.4 % bromophenol blue). The cell extracts were heated for 5 min at 95°C, sonicated for 5 sec at 14 Hz, and loaded in SDS-polyacrylamide gel. The proteins were separated by electrophoresis at 30 mA and then transferred to 0.45 µm Protan nitrocellulose membranes (Amersham). The membranes were blocked for 30 min in 5% milk-PBS and then incubated with primary and the corresponding secondary antibody conjugated to IRDye 680 or IRDye800 (LI-COR). For incubation with streptavidin-IRDye 800, the membrane was blocked and incubated with 1%BSA-PBS. Samples were acquired at Odyssey M Imager (LI-COR Biosciences).

### Phylogenetic tree analysis

The coding sequences (CDS) for rotavirus NSP2 and NSP5 proteins were in silicon translated into amino acids sequences using EMBOSS ‘transeq’ (http://emboss.open-bio.org). The protein sequences were aligned using ‘mafft’ (MAFFT v7.475 (2020/Nov/23) (https://mafft.cbrc.jp/alignment/software/) (73, 74), and the aligned protein sequences were back-translated using EMBOSS ‘tranalign’ (http://emboss.open-bio.org). The phylogeny from the nucleotide multiple sequence alignments were then inferred by using ‘BEAUTi’ and ‘BEAST’ (v1.10.4) (75). In brief, 10’000’000 MCMC steps with the Juke-Cantor model were performed, saving each ten-thousandth tree. After the burn-in of 100,000 states, consensus trees for NSP2 and NSP5, respectively, were calculated and visualized using FigTree (https://beast.community).

## Acknowledgments

The University of Zurich supported this work. The funders had no role in study design, data collection, interpretation, or the decision to submit the work for publication.

## Author contributions

The authors declare no conflict of interest.

Conceptualization: ML, AC, CE; Methodology: ML, AC, CE; Software: KT, CA; Validation: ML, AC, KT, CA, CE; Formal analysis: ML, KT, CA, CF, CE; Investigation: ML, AC, KT, CA, CE; Resources: CE; Data Curation: CE; Writing-Original Draft: ML, CE; Writing-Review & Editing: ML, AC, KT, CA, CF, CE; Visualization: CE; Supervision: CE; Project Administration: CE; Funding Acquisition: CF.

